# Modulation of Secretory Lysosomes During NK Cell Education Leads to Accumulation of Granzyme B and Enhanced Functional Potential

**DOI:** 10.1101/305862

**Authors:** Jodie P. Goodridge, Benedikt Jacobs, Michelle L. Saetersmoen, Dennis Clement, Trevor Clancy, Ellen Skarpen, Andreas Brech, Johannes Landskron, Christian Grimm, Aline Pfefferle, Leonardo Meza-Zepeda, Susanne Lorenz, Merete Thune Wiiger, William E. Louch, Eivind Heggernes Ask, Lisa L. Liu, Vincent Yi Sheng Oei, Una Kjällquist, Sten Linnarsson, Sandip Patel, Kjetil Taskén, Harald Stenmark, Karl-Johan Malmberg

## Abstract

Inhibitory signaling during natural killer (NK) cell education translates into increased responsiveness to activation; however the intracellular mechanism for functional tuning by inhibitory receptors remains unclear. We found that educated NK cells expressing self-MHC specific inhibitory killer cell immunoglobulin-like receptors (KIR) show accumulation of granzyme B, localized in dense-core secretory lysosomes, converged close to the centrosome. This discrete morphological phenotype persists in self-KIR^+^ NK cells independently of transcriptional programs that regulate effector function, metabolism and lysosomal biogenesis. The granzymeB dense, large secretory lysosomes in self-KIR^+^ NK cells were efficiently released upon target cell recognition, contributing to their enhanced cytotoxic capacity. Secretory lysosomes are part of the acidic lysosomal compartment, which has been shown to channel calcium and mediate intracellular signalling in several cell types. Interference of signaling from acidic Ca^2+^ stores in primary NK cells reduced both target-specific Ca^2+^-flux, degranulation and cytokine production. Furthermore, inhibition of PI(3,5)P_2_ synthesis or genetic silencing of the PI(3,5)P_2_-regulated lysosomal Ca^2+^-channel TRPML1 led to increased levels of granzyme B and enhanced functional potential. These results indicate an intrinsic role for lysosomal homeostasis in NK cell education.

## Introduction

Natural killer (NK) cells achieve specificity through unique combinations of variable germ-line encoded receptors. These receptors are critical for the development of cell-intrinsic functional potential, enabling spontaneous activation upon recognition of target cells displaying reduced class I MHC expression.^1^ Inhibitory interactions with self-MHC translate into a predictable quantitative relationship between self-recognition and effector potential, a process termed NK cell education.^2^ Despite being clearly evident in different species,^3^ NK cell education operates through an as yet largely unknown mechanism. Paradoxically, mature NK cells expressing self-MHC specific inhibitory receptors, receiving constitutive inhibitory input during homeostasis, exhibit increased levels of functionality upon ligation of activating receptors.^2, 4, 5, 6, 7, 8, 9, 10^

Mouse models have demonstrated that this functional phenotype is dynamic and dependent on the net signaling input to NK cells during cell-to-cell interactions with both stromal and hematopoietic cells.^11^ Transfer of mature NK cells from one MHC environment to another results in reshaping of the functional potential based on the inhibitory input of the new MHC setting.^12^ Alternatively, genetic knock-down of SLAM-family receptors by CRISPR/Cas9 leads to hyperfunctionality,^10^ whereas deletion of the inhibitory signaling through ITIM and SHP-1 renders NK cells hypofunctional.^4, 13^ However, it remains unclear how and when the net signaling input from activating and inhibitory receptors during NK cell education is integrated to tune the functional potential of the cell. How do NK cells remember their education?

One difficulty in establishing the cellular and molecular mechanisms that account for the calibration of NK cell function is the lack of a steady-state phenotype that defines the educated NK-cell state. Functional readouts used to distinguish self-specific NK cells from hyporesponsive NK cells do not provide information about the prior events that culminate in the development of effector potential. Apart from differences in the relative levels and distribution of NK cell receptors at the cell membrane,^14, 15^ transcriptional and phenotypic readouts at steady state provide scant differences between self and non-self specific NK cells.^16, 17^ Whether inhibitory signaling is converted into a paradoxical gain of function through an as yet unknown mechanism (e.g. arming/stimulatory licensing), or whether expression of self-specific inhibitory receptors protect the cell from tonic activation that would otherwise lead to erosion of function over time (e.g. disarming/inhibitory licensing) remains to be determined.^3, 18, 19^

Here we show that expression of self-specific inhibitory receptors influences the structural organization of the endolysosomal compartment. This allows educated NK cells to sequester granzyme B and mount strong, receptor-triggered effector responses from pre-existing large dense-core secretory lysosomes (also referred to as lytic granules). Moreover, the secretory lysosomes form part of the acidic Ca^2+^ stores in the cells and contribute to the global Ca^2+−^flux and downstream effector function in educated NK cells. These findings connect homeostatic receptor input to lysosomal homeostasis, which tune the functional potential in self-KIR^+^ NK cells.

## Results

### Carrying inhibitory receptors for self HLA class I is associated with increased Granzyme B levels in primary resting NK cells

NK cell education refers to the paradoxical phenomena that NK cells expressing self-specific inhibitory receptors show greater intrinsic functional responsiveness to stimulation through activating receptors.^18^ This intrinsic functional set point determines the response at the single cell and population level in a wide range of read outs and across species.^7^ The impact of NK cell education on degranulation of primary NK cells expressing self versus non-self specific KIR was examined in 96 healthy blood donors (Fig. 1a). In line with several studies over the last decade, NK cells expressing self-specific KIR exhibited greater degranulation in response to HLA class I-deficient K562 cells. To address the mechanisms involved in tuning of effector potential, the expression of cytotoxic effector molecules was monitored in mature NK cells stratified on the expression of self versus non-self specific KIR. The stochastic expression of KIR in NK cells occurs independently of MHC setting, providing a unique situation in which self and non-self specific KIR^+^ subsets can be examined within each individual as a natural equivalent of gene-silencing.^20, 21^ This allowed us to address the impact of reciprocal presence or absence of a self-KIR on the total granzyme B content within equivalent subsets in each individual. Clustering of NK cell phenotypes using t-distributed stochastic neighbor embedding (tSNE) revealed high expression of granzyme B in NK cell subsets expressing self-specific KIR (Fig. 1b). Extended analysis of 64 healthy donors showed significantly higher expression of granzyme B in NK cells positive for KIR2DL3 (2DL3) relative to KIR2DL1 (2DL1) from individuals homozygous for the 2DL3 ligand, HLA-C1/C1. Conversely, granzyme B was elevated in 2DL1^+^ cells from individuals homozygous for the 2DL1 ligand, HLA-C2/C2 (Fig. 1c)., In order to control for the stage of differentiation, which is known to influence expression of effector molecules,^22^ these analyses were performed in NK cells that were NKG2A negative and CD57 negative. Corroborating the link between inhibitory input through self-KIR and granzyme B expression, donors that were heterozygous for HLA-C1/C2 had similarly high levels of granzyme B in both 2DL1 and 2DL3 single-positive NK cells. Granzyme B expression was also higher in 3DL1^+^ NK cells from donors positive for its cognate ligand HLA-Bw4 (Fig. 1d). NK cells with higher levels of 3DL1 surface expression, also known to have a higher functional capacity, ^23^ exhibited greater expression of granzyme B **(Supplementary Fig. 1a)**. It is well established that NKG2A/HLA-E interactions contribute to education of NK cells.^24, 25, 26^ In line with the results of single KIR^+^ NK cell subsets, NKG2A^+^KIR^−^CD57^−^ NK cells expressed slightly higher levels granzyme B **(Supplementary Fig. 1b)**.

**Figure 1.**
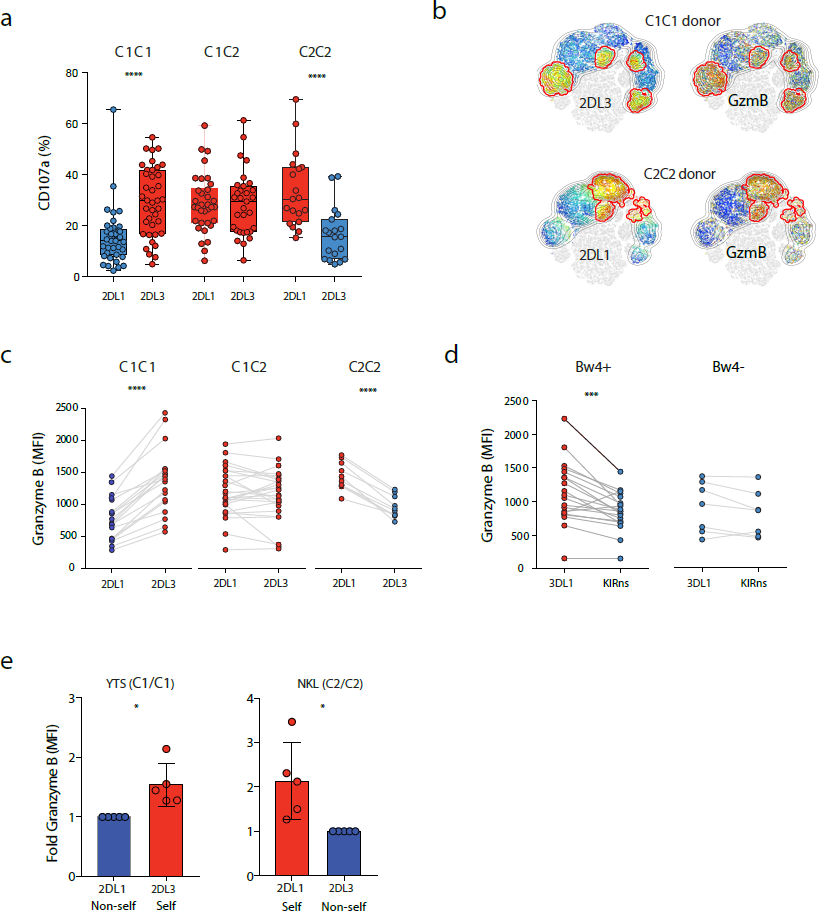
NK cell education through self-KIR is associated with accumulation of granzyme B. (**a**) CD107a expression in the indicated NK cell subset in single-positive NK cells from C1/C1, C1/C2 and C2/C2 donors (n=96). (**b**) SNE plot showing intensity of granzyme B in clusters defined by 2DL3 and 2DL1 expression in C1/C1 and C2/C2 donors, respectively. (n=10) **(c)** Expression of granzyme B in 2DL3 and 2DL1 single-positive NK cells from C1/C1, C1/C2 and C2/C2 donors. **(d)** Expression of granzyme B in 3DL1^+/−^ NK cells from Bw4^+^ (n=20) and Bw4^−^ donors (n=7). **(e)** Expression of granzyme B in YTS and NKL cells transfected with 2DL3 or 2DL1. Graph shows data from one representative experiments.

To study the effect of self KIR expression in a dynamic model, retroviral transduction was used to introduce full length 2DL1 or 2DL3 into NK cell lines YTS (HLAC1/C1) and NKL (HLA-C2/C2). Extending the findings with *ex vivo* staining of primary NK cells, the transduced NK cell lines showed a similar accumulation of granzyme B following transfection of a self KIR (Fig. 1e) and enhanced functionality (**Supplementary Fig. 1c-d**). These data show that the expression of inhibitory self-KIR or NKG2A are connected to the granzyme B load of an NK cell, establishing a link between inhibitory input and the regulation of the core cytolytic machinery of NK cells.

### Inhibitory input during NK cell education influences the granzyme B levels independently of transcriptional cell differentiation programs

To address whether the increased levels of granzyme B in educated NK cells were due to increased transcription, NKG2A^−^CD57^−^ NK cells were sorted by FACS into 2DL3 or 2DL1 single positive populations from C1/C1 and C2/C2 donors and transcriptionally profiled using RNA-Seq. In line with previous studies in mice ^14^, there was a near perfect correlation between genes expressed in self and non-self KIR^+^ human NK cell subsets, including granzyme B and genes encoding transcription factors for effector loci, lysosomal biogenesis, and mechanistic target of rapamycin (mTOR) regulated metabolism (Fig. 2a and **Supplementary Table 1**). For reference we also sorted and performed RNASeq on NK cell subsets at discrete stages of NK cell differentiation CD56^bright^, CD56^dim^NKG2A^−^KIR^−^ and CD56^dim^NKG2A^−^KIR^+^ (**Supplementary Fig. 2**). As expected, NK cell differentiation and KIR acquisition was associated with increased transcription of granzyme B and several other genes known to be involved in regulating effector function, including IRF4 and PRDM1 (**Supplementary Fig. 2**).^27^ These results were confirmed in a panel of 8 selected qPCR targets comprising transcription factors and canonical cell surface markers linked to NK cell differentiation (Fig 2b and Supplementary Fig. 3). Together, these data demonstrated that the increased levels of granzyme B detected by flow cytometry in self-KIR^+^ NK cells occur independently of transcriptionally regulated programs, including differences in tonic metabolic input to the cell.

**Figure 2.**
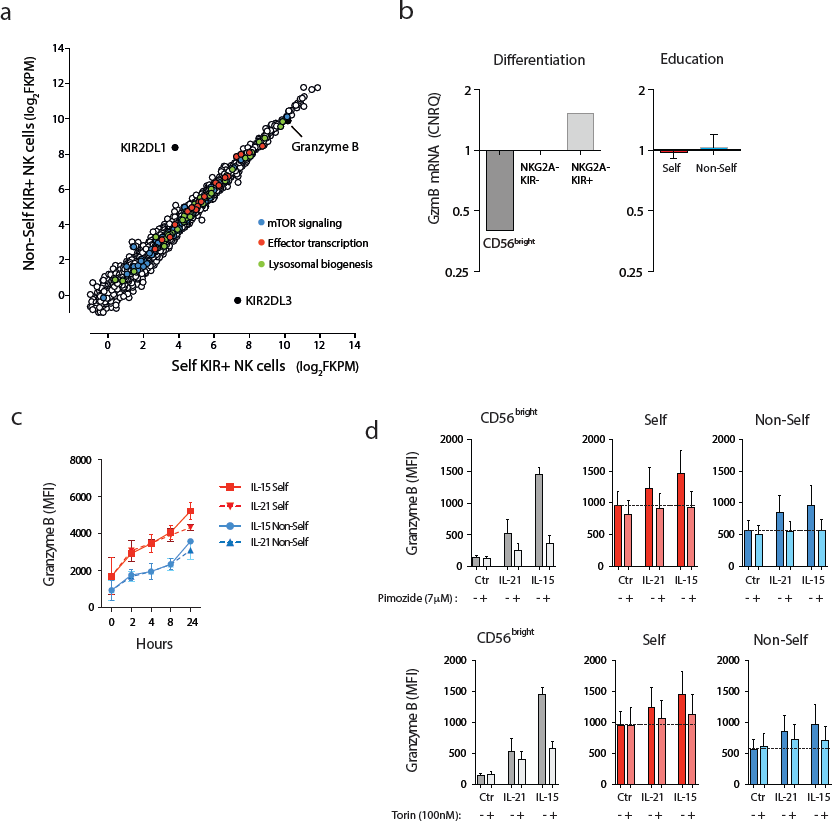
Granzyme B accumulation in educated NK cells is independent of transcriptional programs. **(a)** Global RNA-Seq of sorted single-KIR^+^ NK cells from one representative C1/C1 donor out of three independent donors. **(b)** Quantitative PCR of granzyme B mRNA in in sorted NK cells at different stages of differentiation (left) and in NKG2A-CD57-single-positive NK cells expressing a self- or non-self KIR. (n=5). **(c)** Expression of granzyme B in the indicated NK cell subsets following stimulation with IL-15 or IL-21 for the indicated length of time (n=2). **(d)** Expression of granzyme B after 24h of stimulation with IL-21 or IL-15 in the indicated NK cell subset in the presence or absence of the STAT-5 inhibitor Pimozide (top) (7μM) and the mTOR inhibitor Torin-1 (100nM) (bottom) (n=5).

**Figure 3.**
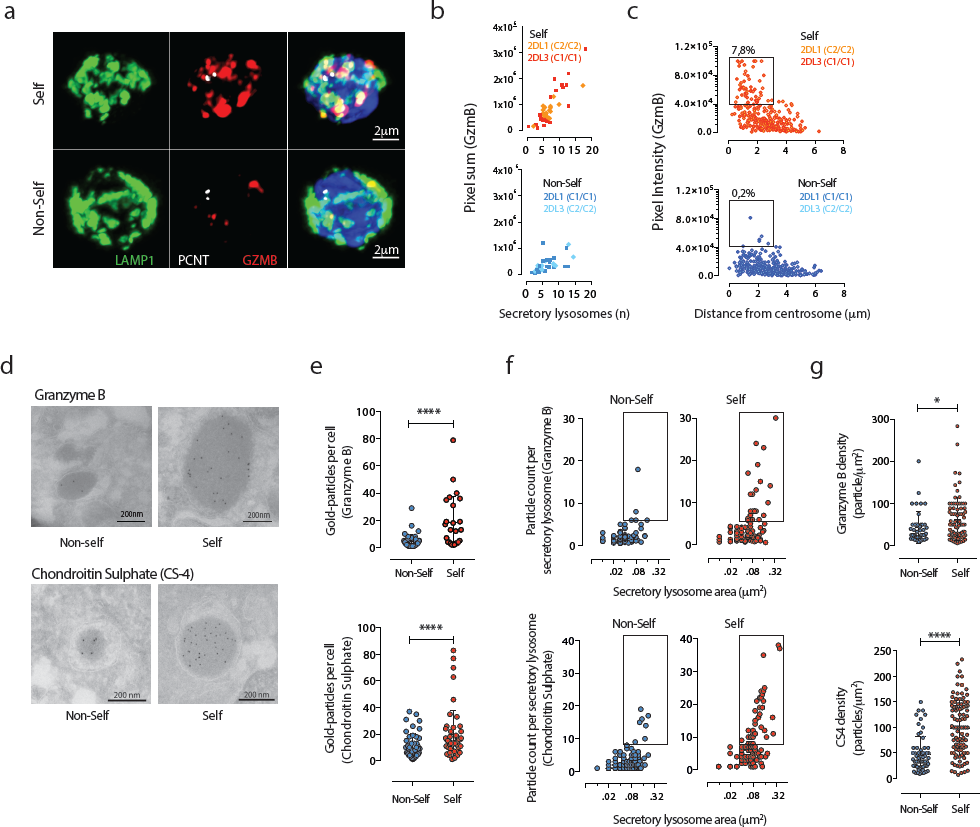
Accumulation of converged dense-core secretory lysosomes in self-KIR^+^ NK cells. **(a)** Confocal microscopy Z-stack showing Pericentrin (PCNT), LAMP-1 and granzyme B (GZMB) staining in sorted CD56^dim^ NKG2A^−^CD57^−^ NK cells expressing Non-Self or Self KIR. **(b)** The pixel sum of granzyme B staining in cells expressing Non-Self (bottom) or Self KIR (top) versus the number of lysosomes. Data are aggregated from sorted 2DL1 and 2DL3 single-positive NK cell subsets from C1C1 (n=5) and C2C2 (n=5) donors. **(c)** Granzyme B expression levels in individual lysosome in NK cells expressing Non-Self or Self KIR versus the distance from the centrosome (804 lysosomes from 3 donors were analysed). Gates were set manually to quantify granzyme-B dense secretory lysosomes **(d)** Representative immuno-EM image showing staining with gold-particle coated anti-granzyme B (top) and chondroitin Sulphate-4 (CS-4) (bottom) of sorted CD56^dim^ NKG2A^−^CD57^−^ NK cells expressing Non-Self or Self KIR. **(e)** Number of gold particles (granzyme B and CS-4) per cell. (Non-Self n=83, Self n=109. Data are from 5 donors and 5 experiments). **(f)** Particle count (granzyme B and CS-4) as a function of the lysosomal area. **(g)** Density of gold particles (granzyme B and CS-4) per lysosomal area (μm^2^).

In mouse NK cells, expression of granzyme B is regulated by cytokine-induced translation from a preexisting pool of mRNA transcript.^28^ Therefore, we explored the possibility that self and non-self specific NK cells may respond differentially to cytokine priming *in vivo,* resulting in divergent steady-state levels of expressed granzyme B. To address this possibility, NK cells exposed to IL-15 or IL-21 for various lengths of time were monitored for granzyme B content using flow cytometry (Fig. 2c). Both self and non-self KIR^+^ CD56^dim^ NK cells displayed increased levels of granzyme B in response to IL-15 and IL-21 stimulation. Notably, the relative differences in granzyme B between self and non-self specific NKG2A^−^CD57^−^ NK cells were similar after stimulation with IL-15 or IL-21 (Fig. 2c). Furthermore, blockade of STAT-5 and mTOR signaling with Pimozide and Torin-1, respectively, abolished the cytokine-induced increase in granzyme B in both self and non-self specific NK cells (Fig. 2d). These data indicate that the observed differences in granzyme B levels at rest are stable and refractory to interference of either STAT or mTOR signaling, suggesting that a stable pool of granzyme B is retained by NK cells independently of constitutive input through cytokine or metabolic signals.

### Accumulation of dense-core secretory lysosomes in self-KIR NK cells

The finding that self-KIR^+^ NK cells expressed higher levels of granzyme B independently of gene expression provided an initial insight into possible post-transcriptional mechanisms underlying the increased functional potential associated with NK cell education. Granzyme B is sequestered into secretory lysosomes with acidic pH.^29^ To determine whether the increased levels of granzyme B in self-specific NK cells were the result of higher density, number, or size of such secretory lysosomes, NKG2A^−^CD57^−^ NK cells were sorted into self- or non-self specific NK cell subsets, imaged by confocal microscopy and analysed in a blinded fashion. Corroborating the difference in granzyme B expression observed using flow cytometry, self-KIR^+^ NK cells had a higher overall intensity of granzyme B staining (Fig. 3a and **Supplementary Fig. 4a**). Self-KIR^+^ NK cells displayed an increased level of fluorescence intensity for granzyme B (Fig. 3b), which in turn correlated with proximity to the centrosome (Fig. 3c). These granzyme-B dense secretory lysosomes, localised close to the centrosome were uniquely found in educated self-KIR^+^ NK cells. Blinded scoring of the size and staining intensity of 804 granzyme-B^+^ structures (secretory lysosomes) from 20 single-KIR^+^ NK cells, revealed that 50% of the educated NK cells carried at least one large secretory lysosome (3,2 on average), representing on average 7,8% of the total number of secretory lysosomes in the cell. Notably, in these cells, such granzyme-B dense lysosomes contributed to 36% (8-76%) of the total granzyme B signal.

**Figure 4.**
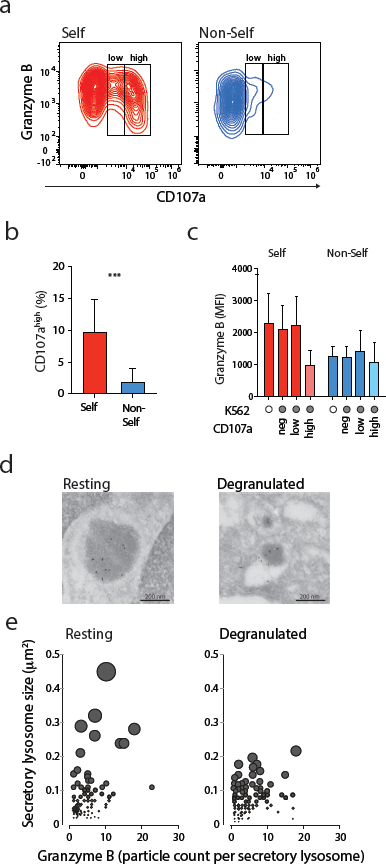
Self-KIR^+^ NK cells mobilize large dense-core secretory lysosomes and release granzyme B down to base-line levels following target cell stimulation. **(a)** Representative example of granzyme B and CD107a expression in Self KIR+ and Non-Self KIR^+^ CD56^dim^ NKG2A^−^CD57^−^ NK cells following stimulation with K562 cells. **(b)** Aggregated data of percent CD107a^high^ NK cells following stimulation with K562 cells (n=5). **(c)** Expression of granzyme B in the indicated NK cell subset after stimulation with K562 (n=4 C1/C1 donors). **(d)** Representative Immuno-EM image of resting or sorted CD107a^high^ NK cells. **(e)** Secretory lysosome size and granzyme B content as determined by immuno-EM in resting and sorted CD107a^high^ NK cells after stimulation with K562.

Optical resolution limits of confocal microscopy prevented accurate assessment of organelle size. To more precisely analyse areas with high granzyme B intensity and density of individual secretory lysosomes, sorted self-KIR^+^ and non-self KIR^+^ NK cells were sectioned and stained for immuno-electron microscopy (Immuno-EM) using anti-granzyme B mAb and Protein A-gold (Fig. 3d). Quantification of gold particles per cellular section revealed overall greater granzyme B staining in self-KIR^+^ NK cells, consistent with both the flow cytometry and confocal microscopy (Fig. 3d-e). Since retention of granzyme B and re-loading of secretory lysosomes depends on the serglycin content in the matrix of the secretory lysosomes,^30^ sections of self-KIR^+^ and non-self KIR^+^ NK cells were stained for expression of Chondroitin Sulphate 4 (CS4), a predominant glycosaminoglycan side-chain associated with serglycin in cytotoxic lymphocytes.^31^ Self-KIR^+^ NK cells had a higher overall intensity of CS4-staining (Fig. 3d-e), suggesting that NK cell education leads to changes in the matrix composition of the secretory lysosomes. Granzyme B and CS4 staining further revealed a small increase in the average secretory lysosome size, resulting in larger total secretory lysosomal areas, without significant difference in relative cell size (**Supplementary Fig. 4b-d**). Similarly, analysis of gold particle distribution against secretory lysosome area in immuno-EM images revealed that self-specific NK cells had larger granzyme B-dense lysosomal areas (Fig. 3f), and overall greater secretory lysosome densities (Fig. 3g). These data provide a link between the expression of self-specific inhibitory KIR and retention of enlarged, granzyme B-dense secretory lysosomes.

### Self-KIR^+^ NK cells mobilize large dense-core secretory lysosomes and release granzyme B to base-line levels following target cell stimulation

It is well established that self-KIR^+^ NK cells display stronger degranulation responses than non-self KIR^+^ NK cells at the population level,^2^ which was corroborated in our functional analysis of 96 donors (Fig. 1a). Furthermore, recent studies suggest that release of as few as one secretory lysosome can lead to target cell killing.^32^ Given the unique accumulation of granzyme B-dense secretory lysosomes in self-KIR^+^ NK cells we examined their fate following stimulation with K562 cells. Granzyme B release in self KIR^+^ NK cells was associated with strong mobilization of secretory lysosomes, reflected in a higher discrete mode of CD107a expression (CD107a^high^) (Fig. 4a-b). High CD107a expression was accompanied by a decrease in granzyme B expression to the levels observed in non-self specific KIR^+^ NK cells (Fig. 4c). Immuno-EM of sorted CD107a^high^ NK cells revealed a specific reduction of large and granzyme B-dense lysosomes following target cell interaction (Fig. 4d-e). Thus, the accumulation of dense-core secretory lysosomes during education and their effective release upon stimulation, provide a plausible explanation for the enhanced cytotoxic potential of self-KIR^+^ NK cells and their ability to perform serial killing.^33^

### Compromising lysosomal activity decreases functional potential in NK cells

NK cell education has a global influence on the function of self-KIR^+^ NK cells extending beyond degranulation responses. These include increased Ca^2+^ flux following receptor ligation, increased ability to form stable conjugates and increased cytokine production following target cell interaction.^18, 34^ Therefore, we addressed whether there could be link between the observed morphological changes in the lysosomal compartment and the known enhanced global responsiveness associated with NK cell education. The secretory lysosome has a low pH and belongs to the acidic compartment. There is increasing evidence suggesting that local Ca^2+^ signaling from the acidic compartment, including secretory lysosomes contributes to the spatiotemporal coordination of signaling cascades and boost Ca^2+^ signaling from the endoplasmatic reticulum (ER).^35, 36, 37, 38^ While most of the studies in this area have so far been performed in non-immune cells, there is evidence in both T cells and NK cells, that lysosomal Ca^2+^ release may play an important role in degranulation.^35, 39^ Therefore, we examined whether primary human NK cells were affected by regulation of lysosomal activity and whether this had consequences on their global responsiveness to receptor ligation.

Functional responses of primary NK cells were determined in the presence and absence of glycyl-L-phenylalanine-beta-naphthylamide (GPN), a dipeptide substrate of cathepsin C associated with release of Ca^2+^ from the lysosomes.^40^ GPN causes osmotic permeablization of cathepsin C-positive compartments,^41^ resulting in the collapse of the pH gradient and controlled equilibration of small solutes (dictated by donnan equilibrium), including Ca^2+^, between the acidic compartment and the cytosol.^40^ Treatment of resting primary NK cells with GPN dampened the detection of global Ca^2+^-flux in the cytosol in response to ligation with CD16 or a combination of DNAM-1 and 2B4 (Fig. 5a). GPN treatment alone resulted in a low level of mobilization of CD107a^+^ vesicles to the cell surface (Fig. 5b) but abrogated degranulation and more importantly abrogated the production of IFN-γ in response to K562 cells (Fig. 5b-c). Similar results were obtained using mefloquine, another lysosomotropic agent that specifically disrupts lysosomal homeostasis through buffering of the acidic pH gradient (Fig. 5d).^42, 43^ Thus, disruption of the lysosomal compartment not only affects the mobilization of secretory lysosomes but also the production of cytokines, suggesting that cytokine production in response to stimuli requires lysosomal-derived signals. Importantly, none of these compounds showed any general cellular toxicity at the doses tested as compared to the positive control LLeucyl-L-leucine methyl ester (LeuLeuOMe), a lysosomotropic agent known to induce apoptosis in immune cells through induced lysis of secretory lysosomes (**Supplementary Fig. 5**). Furthermore, GPN treatment did not interfere with degranulation in response to PMA/Ionomycin, which raises cytosolic free Ca^2+^ by directly accessing both intra- and extra-cellular free Ca^2+^ (**Supplementary Fig. 6**). This opens the possibility that differential signaling in self-KIR+ and nonself-KIR+ NK cells may be influenced by lysosomal-derived Ca^2+^ signals.

**Figure 5.**
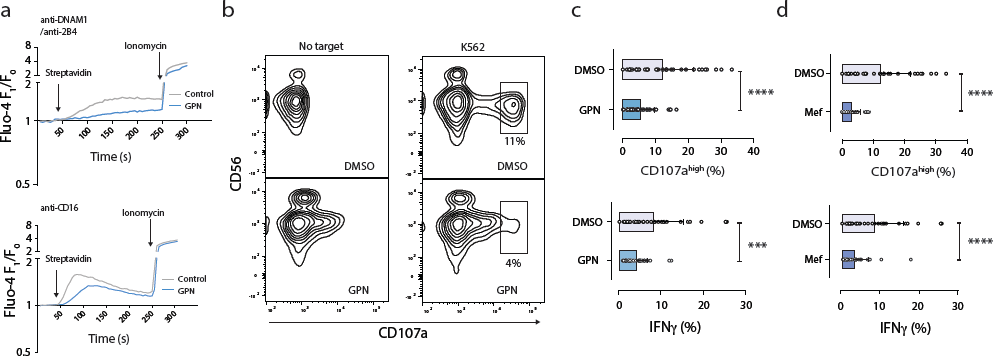
Compromising lysosomal activity decreases functional potential in NK cells. **(a)** Global Ca^2+^-flux measured by Fluo-4 F_1_/F_0_ ratio following stimulation with biotinylated anti-DNAM-1/anti-2B4 (top) or biotinylated anti-CD16 (bottom) crosslinked at the indicated time-point with streptavidin in the presence (added before stimulation and maintained throughout the assay) or absence of GPN (50μM). **(b)** Representative example of granzyme B and CD107a expression following stimulation of NK cells with K562 cells in the presence or absence of GPN. Frequency of CD107^high+^ (top) and IFN-γ^+^ (bottom) NK cells following stimulation with K562 cells in the presence or absence of **(c)** 50uM GPN (n=32) and **(d)** 10uM Mefloquine (n=32).

**Figure 6.**
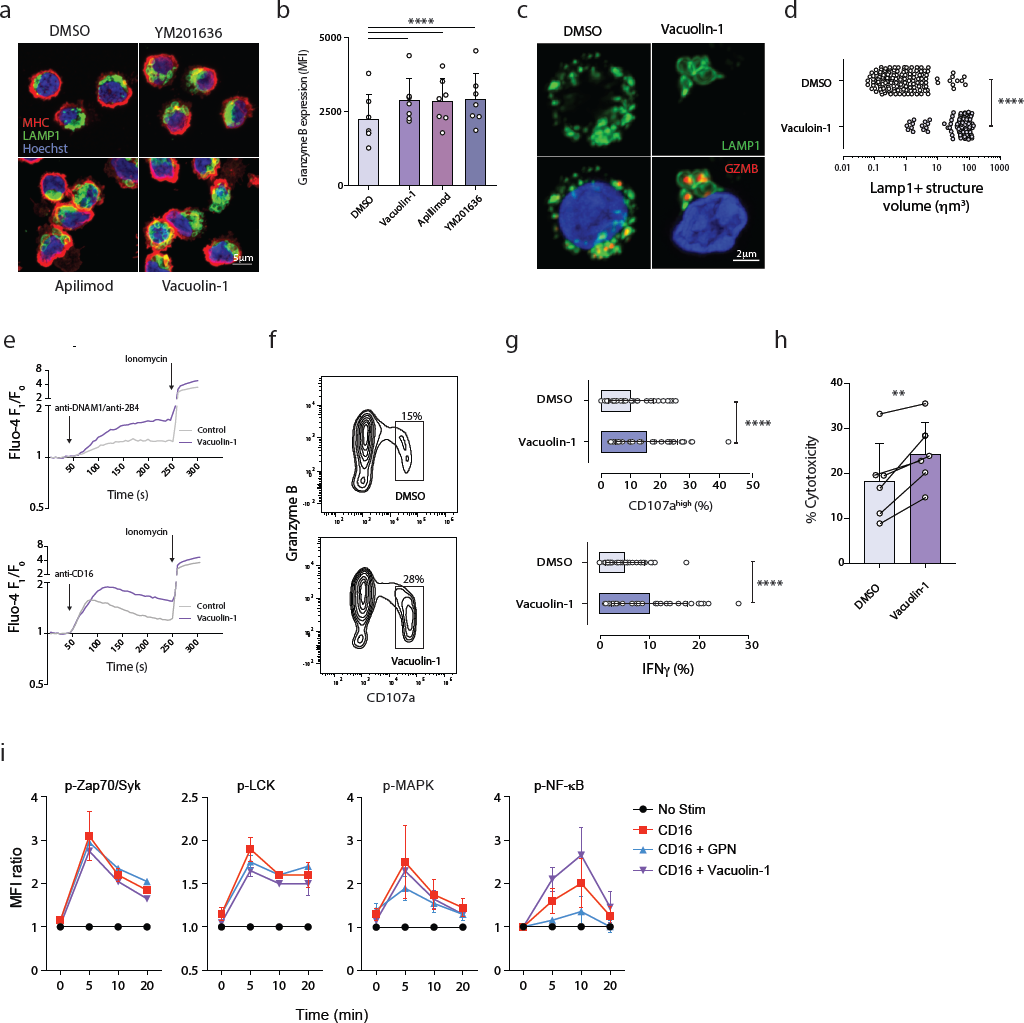
Enlarging the secretory lysosomes leads to enhanced NK cell functionality. **(a)** Confocal Z-stack showing MHC-I, LAMP-1 and granzyme B (GZMB) staining in primary NK cells following PIKfyve inhibition using overnight incubation with 1uM vacuolin-1, 1uM Apillimod or 1uM YM201636. **(b)** Intracellular granzyme B expression following incubation with the indicated PIKfyve inhibitor assessed by flow cytometry (n=7). **(c)** Representative example of confocal image of primary NK cells treated overnight with 1uM vacuolin-1 or DMSO. **(d)** Compiled confocal data on the volume of LAMP-1^+^ structures from cells treated with DMSO or vacuolin-1 as in c (n=149). **(e)** Ca^2+^-flux in NK cells in response to stimulation with biotinylated anti-DNAM-/2B4 (top) or anti-CD16 (bottom) crosslinked with streptavidin at the indicated timepoint. Cells were treated with 10uM vacuolin-1 added directly before the assay and then maintained through the incubation time. **(f)** Representative FACS histogram of granzyme B versus degranulation following stimulation with K562 in the presence of DMSO or 10μM vacuolin-1. **(g)** Frequency of CD107a^high+^ (top) and IFN-γ^+^ N K cells (bottom) after stimulation with K562 in the presence of DMSO or 10μM vacuolin-1. **(h)** FACS-based killing assay showing NK cell killing of K562 cells after treatment with DMSO or 10μM vaculin-1 (n=6) **(i)** Relative phosphorylation of the indicated signaling molecules following stimulation with biotinylated anti-CD16 (10ug/mL) crosslinked with avidin in the presence of 50μM GPN or 10μM vacuolin-1.

### Enlarging the secretory lysosomes leads to enhanced NK cell functionality

The lysosomal compartment undergoes constant modulation through Ca^2+^ regulated fission and fusion events.^44, 45^ Lysosomal fission is dependent on Ca^2+^ release via a lysosome-specific channel, transient receptor potential mucolipin-1 (TRPML1).^46, 47, 48^ TRPML1 is activated by phosphoinositide 3,5-bisphosphate PI(3,5)P_2,_^49^ and may prevent uncontrolled fusion with secretory lysosomes.^50^ To probe this axis, we blocked PI(3,5)P_2_ synthesis in resting NK cells using three chemically distinct small molecule inhibitors of the PI3P 5 kinase, PIKfyve.

Treatment of cells with vacuolin-1, apilimod and YM201636,^51^ led to enlargement of the lysosomal compartment (Fig. 6a). Importantly, this was accompanied by a modest increase in granzyme B levels (Fig. 6b). Confocal microscopy of vacuolin-1 treated primary NK cells revealed localization of granzyme B within enlarged LAMP-1^+^ structures (Fig. 6c-d). Treatment of NK cells with vacuolin-1 also increased global Ca^2+^ flux in response to receptor-ligation (Fig. 6e) and enhanced specific degranulation (and mobilization of granzyme B) and IFNγ production in response to stimulation by K562 cells (Fig. 6f-g). Furthermore, the increased granzyme B expression and degranulation following PIKfyve inhibition correlated with increased natural cytotoxicity against K562 cells (Fig. 6h). These results demonstrate that chemical blockade of PIKfyve results in enlargement of the lysosomal compartment and enhanced NK cell functionality.

In order to identify the point at which lysosomal disruption (GPN) or lysosomal enlargement (vacuolin-1) interfered with intracellular signaling pathways in NK cells, we probed signaling both proximal and distal to the plasma membrane. Vacuolin-1 had a minimal effect on upstream signaling, including ZAP70 and Lck following ligation of CD16 (Fig. 6i). However, the propagation of down-stream signals through NF-κB was increased by treatment with vacuolin-1. Conversely, disruption of lysosomal Ca^2+^-flux by GPN had the reverse effect on CD16-induced NF-κB signaling (Fig. 6i). Hence, physical modulation of the acidic Ca^2+^ stores affects downstream signaling in response to receptor ligation and tunes NK cell effector responses.

### TRPML1-mediated modulation of secretory lysosomes in NK cells

PIKfyve is recruited to PI3P positive compartments where it activates the lysosomal calcium channel TRPML1 via the production of PI(3,5)P_2._^47, 48^ Analysis of the transcriptional levels of TRPML1 in discrete NK cell subsets revealed TRPML1 mRNA was expressed at equal levels in all NK cell subset (Fig. 7a). Agonistic stimulation of TRPML using the chemical compound MK6-83, which activates TRPML1 and TRPML3,^52, 53, 54^ resulted in loss of granzyme B (Fig. 7b) and decreased specific degranulation and IFNγ responses to K562 cells (Fig. 7c). TRPML3 mRNA was not expressed in resting human NK cells (Data not shown). Conversely, silencing of TRPML1 by siRNA in resting primary NK cells led to increased levels of granzyme B (Fig. 7d-e). Moreover, in concordance with the effects of pharmacological inhibition of PIKfyve, siRNA silencing of TRPML1 (Fig. 7f) led to enhanced degranulation and IFN-γ production in primary resting NK cells (Fig. 7g). These results demonstrate a role for TRPML1 in the modulation of granzyme B content and in tuning of effector function in NK cells.

**Figure 7.**
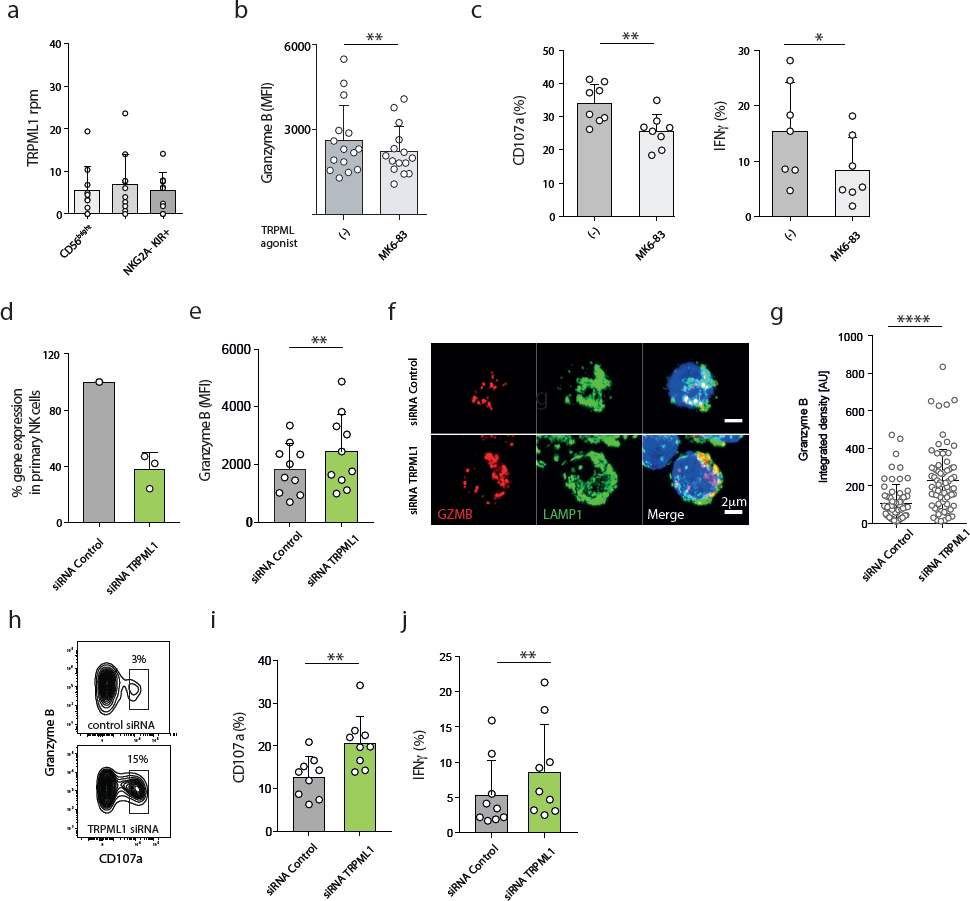
TRPML1-mediated modulation of secretory lysosomes in NK cells. **(a)** mRNA expression (RNA-Seq) of TRPML1 in the indicated NK cell subsets sorted from PBMC and analysed directly. **(b)** Granzyme B expression in NK cells treated for 2 hours with 10μM of the TRPML1 agonist MK6-83. **(c)** Degranulation (left) and IFNγ responses (right) by resting primary NK cells following stimulation with K562 cells for 4 hours in the presence or absence of 10μM MK6-83. **(d)** Relative mRNA expression (qPCR) of TRPML1 after siRNA silencing in resting NK cells. **(e)** Granzyme B expression in NK cells cultured in 1ng/ml of IL-15 for 72 hours after gene silencing by TRPML1 siRNA. **(f)** Confocal microscopy image showing LAMP-1 and granzyme B (GZMB) staining in siRNA TRPML1 silenced NK cells **(g)** Summary of integrated granzyme B intensity per cell as quantified with imageJ (n=17-37 cells from 3 experiments). AU = arbitrary units. **(h)** Representative example of FACS plot showing granzyme B expression versus CD107a in siRNA TRPML1 silenced primary NK cells after stimulation with K562 cells. Compiled data on (**i**) degranulation and (**j**) IFNγ production in TRPML1-silenced primary NK cells. Data are from two independent experiments and three donors with confirmed siRNA silencing.

## Discussion

NK cell education is a dynamic process during which NK cells calibrate their functional potential to self-MHC. However, it has been unclear how receptor input during NK cell education is integrated and retained in order for NK cells to remain self-tolerant, whilst also able to deliver spontaneous, well-tuned functional responses upon subsequent challenges. Our results suggest that unopposed activation signals lead to physical disarming of NK cells, reflected in the TRPML1-induced modulation of the lysosomal compartment. The accumulation of dense-core secretory lysosomes under the influence of inhibitory self-MHC interactions provides mechanistic insights into the paradox of how inhibitory signaling is translated into a state of enhanced functional potential that persists between successive cell-to-cell contacts. The structural change in the lysosomal compartment and loading of dense-core secretory lysosomes may represent a form of molecular memory of receptor signaling during NK cell education.

A variety of models and nomenclature have been used to describe the process of NK cell education. However, regardless of whether the functional phenotype is caused by gain of function (arming/stimulatory licensing) in self-KIR^+^ NK cells or loss of function (e.g. disarming/inhibitory licensing) in nonself-KIR^+^ NK cells ^3, 55^, the net outcome of these processes is a consistent difference in the intrinsic functional potential of cells carrying self- and non-self receptors at rest. A structural basis for the difference in functional responsiveness has recently been proposed,^14, 15^ whereby educated NK cells display a unique compartmentalization of activating and inhibitory receptors at the nano-scale level on the plasma membrane. Complementing this preexisting phenotype, we show that NK cell education is also tightly linked to the accumulation of large, granzyme B-rich secretory lysosomes, located closer to the centrosome in resting NK cells.

An increasing body of evidence supports the role of the acidic compartment not only in the triggering of Ca^2+^ signaling, but also in the spatiotemporal coordination of signaling cascades,^35, 36, 37, 38^ and the regulation of receptor degradation.^56^ In both T cells and NK cells, lysosomal Ca^2+^ release plays an important role in degranulation.^35, 39^ Signaling from the lysosomal compartment was also recently shown to regulate the migratory behavior of dendritic cells in a TRPML1-dependent fashion.^57^ Our data suggest that there is a quantitative relationship between modulation of the lysosomal compartment under the influence of inhibitory receptors and intrinsic functional potential of NK cells. While the exact role of the acidic Ca^2+^ store for the enhanced functional potential in self-KIR^+^ NK cells remains elusive, pharmacological inhibition of Ca^2+^ release from the intracellular acidic stores, together with analysis of Ca^2+^-flux, consistently pointed to a role for the secretory lysosome in propagating surface receptor signaling. Notably, a correlation between the size of the lysosomes and level of Ca^2+^-flux has previously been describe in fibroblasts from patients with Parkinson disease.^58^ On that note, it is tempting to speculate that the gain of natural cytotoxicity in lL15 stimulated CD56^bright^ NK cells may be likewise related to the associated emergence of secretory lysosomes.^59^

We examined the molecular pathway that led to accumulation of secretory lysosomes in self-KIR^+^ NK cells, or rather the lack of accumulation of such lysosomes in non-self KIR^+^ NK cells. The difference in secretory lysosome size and densities in the absence of active lysosomal biogenesis led us to explore the pathways involved in the continuous modulation of the lysosomal compartment through fission and fusion events.^45^ Several activating receptors have been implicated in NK cell education, including NKG2D, SLAM family receptors and DNAM-1.^10, 16, 60^ NKG2D and DNAM-1 signaling activates the PI3K/AKT pathway.^61, 62^ Engagement of inhibitory receptors and SHP-1 signaling block NK cell activation at an early stage of the activation signaling pathway, preventing actin cytoskeletal rearrangement and the recruitment and phosphorylation of activation receptors.^63, 64, 65^ The PI3K/AKT pathway is also controlled by SHIP1, which has also been implicated in tuning the effector function of NK cells during education.^66, 67^ Notably, PIKfyve and TRPML1 are activated downstream of the PI3K/AKT pathway.^56^ Mutations in TRPML1 cause mucolipidosis type IV, which is characterized by enlarged lysosomes.^68, 69^ We hypothesized that continuous unopposed signaling through activating receptors during homeostatic cell-cell interactions may trigger the AKT/PIKfyve/TRPML1 axis and thereby promote lysosomal fission, leading to an inability to sequester granzyme B in large secretory lysosomes (Fig. 8). To explore this hypothesis, we used a combination of pharmacological agonists and antagonists combined with genetic approaches to interfere with the PIKfyve/TRPML-1 pathway. Pharmacological inhibition of PIKfyve by small chemical compounds, including vacuolin-1 and apilimod, is known to cause enlarged lysosomes in several cell types, including mast cells and macrophages.^51, 70, 71^ Here, we show that inhibition of PIKfyve by three different chemical compounds caused enlargement of the lysosomal compartment. Importantly, this was associated with increased granzyme B expression, increased Ca^2+^-flux, and more potent effector function. A similar functional phenotype was obtained when silencing the lysosome-specific Ca^2+^ release channel TRPML1. Together these data support a model where tonic or intermittent activation signals through the PI3K/AKT pathway result in PIKfyve activation and TRPML1-induced lysosomal fission, ultimately leading to lack of large secretory lysosomes and reduced functional potential in NK cells.

**Figure 8.**
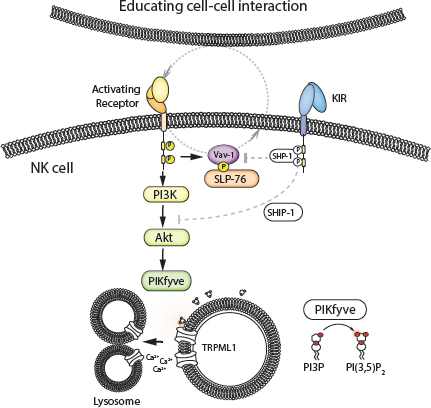
Model describing the putative pathway downstream of activating receptors leading to activation of PIKfyve and TRPML1, lysosomal fission and a hyporesponsive state. Self-specific inhibitory KIRs interfere with activation signals at a proximal level and allow accumulation of dense-core secretory lysosomes.

In Chediak-Higashi Syndrome (CHS), mutation of the *LYST* gene leads to the formation of giant secretory lysosomal structures.^72^ NK cells in CHS patients are hyperresponsive and hypersecretory but are unable to degranulate.^73, 74^ Although NK cell activation followed by secretory lysosome convergence and polarization appears to be normal in *LYST*-deficient NK cells, the enlarged secretory lysosomes fail to pass through the cortical actin meshwork openings at the immunological synapse.^75^ By monitoring lysosomal size and granzyme B density prior to and following degranulation, we observed a selective loss of the pre-converged, large secretory lysosomes after degranulation. These large lysosomes had an area above 0,2 μm^2^, corresponding to a diameter of around 500nm, pointing to a possible difference in the density of the actin meshwork and physical restriction of degranulation between resting primary NK cells and NK92 cells. However, another contributing factor may be LYST-mediated modulation of lysosomal size during NK cell activation allowing smaller proportions of the large lysosomes to be released during the effector response, as has been demonstrated for degranulation in mast cells.^76^

An outstanding question is how lysosomal fission leads to loss of matrix component and lower levels of granzyme B in non-self KIR^+^ NK cells. Granzyme B can be synthesized and secreted directly through the constitutive secretory pathway.^77^ It is possible that an enhanced rate of lysosomal fission during weakly agonistic cell-cell interactions and the corresponding failure to accumulate dense-core lysosomes in NK cells lacking self-specific receptors leads to loss of granzyme through the secretory route.^78^ Indeed, NK cells in serglycin^−/−^ mice lack dense-core lysosomes, retain less granzyme B which is secreted from the cell at a greater rate, and exhibit reduced degranulation in response to stimuli.^30^

Another remaining challenge is to decipher when and where TRPML1-mediated physical disarming takes place. Transfer experiments in mice have established an indisputable role for cell-to-cell interactions in shaping the functionality of mature NK cells.^5, 79, 80^ Although the detailed time-scale and spatial aspects of such cell interactions remain largely unknown, transfer of functional NK cells to MHC-deficient environments leads to induction of hyporesponsiveness.^11^ SHP-1 intersects signaling of activating receptors upstream of Vav-1,^65^ and rapidly shuts down the process of forming an activating NK cell synapse with target cells.^81^ While the inhibitory synapse and the productive cytolytic synapse have been studied in great detail, much less is known about immune synapses formed between resting immune cells during homeostasis. It is possible that cells lacking self-specific inhibitory receptors form a succession of non-cytolytic immune synapses under homeostasis leading to loss of dense-core secretory lysosomes and “leakage” of their functional potential.^78^ It has previously been shown that trans-presentation of IL-15 to NK cells, resulting in activation of AKT is negatively regulated by inhibitory interactions with self MHC.^82^ Thus, it is possible that unopposed constitutive IL-15 activation may occur in NK cells that lack self-specific inhibitory KIR, which in turn would affect lysosome stability and/or retention through the mechanism described here.

The dose dependent induction of granzyme B expression in response to cytokine, or viral infection, is connected to activation of the metabolic check-point kinase mTOR.^83^ Notably, however, we did not observe any transcriptional imprint in the mTOR pathway when we examined circulating NK cells at rest, arguing against a major role for metabolism in the persistence of the distinct organization of the lysosomal compartment seen in circulating blood self-KIR^+^ NK cells. It was recently shown that mTOR activation contributed to the functional rheostat during effector responses in educated murine NK cells.^84^ Interestingly, mTOR activation and function are dependent on its lysosomal localization and the vacuolar H(+) -ATPase (VATPase) activity.^85^ Furthermore, TRPML1 provides a negative feedback loop on mTOR activity.^50^ Thus, the difference in lysosomal composition described in the present study could potentially contribute to enhanced mTOR activation observed in educated NK cells upon stimulation.^84^

In conclusion, our findings suggest a mechanism by which NK cell education operates through modulation of the lysosomal compartment under the influence of inhibitory receptor ligand interactions. Qualitative differences in the morphology of the lysosomal compartment and signaling from acidic Ca^2+^ stores, allow the cytolytic machinery to operate independently of transcription during the effector response. Furthermore, the data suggest that it may be possible to boost NK cell functionality through targeted manipulation of Ca^2+^ homeostasis within lysosome-related organelles.

## Methods

### Cells

Buffy coats from random healthy blood donors were obtained from the Karolinska University Hospital and Oslo University Hospital Blood banks with informed consent. Peripheral blood mononuclear cells were separated from buffy coats by density gravity centrifugation (Lymphoprep; Axis-Shield) using fretted spin tubes (Sepmate; Stemcell Technologies). Genomic DNA was isolated from 200µl of whole blood using DNeasy Blood and Tissue Kit (Qiagen). KIR ligands were determined using the KIR HLA ligand kit (Olerup SSP) for detection of the HLA-Bw4, HLA-C1, and HLA-C2 motifs. NK cells were purified using negative selection (Miltenyi) with an AutoMACS Pro Seperator. K562 cells were maintained in RPMI +10%FCS.

### Phenotyping by Flow Cytometry

Isolated PBMC were stained for flow cytometric analysis using an appropriate combination of antibodies as detailed in the supplementary experimental procedures. After surface staining, cells were fixed and permeabilized using a fixation/permeabilization kit (BD Bioscience Cytofix/Cytoperm) prior to intracellular staining with anti-granzyme B-A700 (GB11). Samples were acquired using an LSRII flow cytometer (Becton Dickinson) and data was analyzed using FlowJo V10.0.8 (TreeStar). Stochastic neighbor embedding (SNE) analysis was performed as described in supplementary experimental procedures

### RNAseq and qPCR

RNASeq was performed using single-cell tagged reverse transcription (STRT), a highly multiplexed method for single-cell RNA-seq. Real time quantitative PCR was used to study the difference in the expression of 20 genes of interest in sorted differentiation and education subsets of NK cells, as detailed in supplementary experimental procedures.

### Confocal fluorescence microscopy and Image analysis

Sorted NK cells were prepared for confocal microscopy using fixation/permeabilization (BD Bioscience Cytofix/Cytoperm) prior to intracellular staining with mouse anti-human granzyme B-A647 (GB11), rabbit anti-human pericentrin (Ab4448) followed by Donkey anti-Rabbit IgG Alexa555. After staining, the fixed cells were adhered to glass cover slips using Cell Tak (Corning) and mounted using Pro-long Gold Antifade with DAPI. The cells were examined with a Zeiss LSM 710 confocal microscope (Carl Zeiss MicroImaging GmbH, Jena, Germany) equipped with an Ar-Laser Multiline (458/488/514nm), a DPSS-561 10 (561nm), a Laser diode 405-30 CW (405nm), and a HeNe-laser (633nm). The objective used was a Zeiss plan-Apochromat 63xNA/1.4 oil DICII. Image processing and analysis were performed with basic software ZEN 2011 (Carl Zeiss MicroImaging GmbH, Jena, Germany) and Imaris 7.7.2 (Bitplane AG, Zürich, Switzerland). Confocal z-stacks were deconvolved using Huygens Essential 14.06 (Scientific Volume Imaging b.v., VB Hilversum, The Netherlands). ImarisCell was used to identify secretory lysosomes and centrosomes in confocal Z-stacks of single cells, while Imaris Venture was used to find the correlation between the intensity of the individual secretory lysosome and their distance to the centrosome center.

### Electron Microscopy

Sorted NK cells for immuno-EM were fixed in a mixture of 4% formaldehyde and 0.1% glutaraldehyde in 0.1 M PHEM buffer (60 mM PIPES, 25 mM HEPES, 10 mM EGTA and 2 mM MgCl2 at pH 6.9), followed by embedding in 10% gelatin, infiltration with 2.3 M sucrose and frozen in liquid nitrogen (LN2). Ultrathin sections (70-90 nm) of cell pellets were cut on a Leica Ultracut (equipped with UFC cryochamber) at −110C, picked up with a 50:50 mixture of 2.3 M sucrose and 2% methyl cellulose. Sections were then labeled with antibodies against granzyme B (496B, eBioscience) or Chondroitin Sulphate 4 (2B6, AMSBIO), followed by a bridging rabbit-anti-mouse antibody (DAKO, Denmark) and protein A gold (University Medical Center, Utrecht, Netherlands). Microscopy was done at 80 kV in a JEOL_JEM1230 and images acquired with a Morada camera. Further image processing was done in Adobe Photoshop. Quantification was done according to established stereological procedures.

### Functional Assays

Functional assays were performed at 37°C in complete medium (RPMI +10%FCS) for the times indicated. Purified NK cells were incubated with K562 target cells for 5 hours at a ratio of 1:1 in the presence of anti-CD107a-alexa488 (H4A3, Biolegend) for degranulation assays, or with the addition of Brefeldin A (GolgiPlug BD) for degranulation plus intracellular cytokine assays. Lysosomotropic reagents were added prior to addition of targets/stimulation by agonistic antibodies and kept for the duration of the assay using the following final concentrations; Glycyl-LPhenylalanine-β-Naphthylamide (GPN, 50uM), mefloquine (10uM), vacuolin-1 (1-10uM). The TRPML1/3 agonist MK6-83 was used at 10uM.

### Phospho-flow cytometry

Functional assays for phospho-flow cytometry were performed at 37°C in complete medium in NK cell suspensions between 5-10 M/mL for 20 min. Cells were pretreated for 1h using GPN (50uM) or Vacuolin-1 (10uM), after which biotinylated CD16 (Biolegend, clone 3G8) was added to final concentrations of 5 µg/mL each. After 1 min, the aliquot for the 0 min (unstimulated) sample was taken out and mixed with Fix Buffer I (BD Biosciences). After one additional minute, the stimulation was started by crosslinking the biotinylated antibodies with 50 µg/mL avidin (Thermo Fischer Scientific) and the aliquots for the 5 min, 10 min and 20 min samples were transferred into Fix Buffer I (BD bioscience) at the corresponding time points. Cells were fixed at 37°C for 10 min, washed and re-suspended in PBS. To allow combination of the differently stimulated samples into one, two dimensional fluorescent cell barcoding (FCB) was utilized. Samples were stained in distinct concentrations of amine-reactive pacific blue succinimidyl ester (Thermo Fisher Scientific) for the time points (0 min – 0.69 ng/mL, 5 min – 6.25 ng/mL, 10 min – 25 ng/mL and 20 min – 100 ng/mL) in combination with amine-reactive pacific orange succinimidyl ester (Thermo Fisher Scientific) for the different stimulations (control – 10 ng/mL, GPN – 100 ng/mL and Vacuolin-1 – 500 ng/mL). After 20 min at RT, samples were washed twice in wash solution (PBS supplemented with 1% FCS and 0.09% sodium azide), combined, permeabilized (Perm Buffer III, BD Biosciences) and stored at −80°C. For thawing, samples were incubated 20 min on ice. Then, they were washed in wash solution and stained with Alexa Fluor 647-conjugated phospho epitope-specific antibodies against ZAP70/syk (pY319/pY352), Lck (pY505), Erk1/2 (pT202/pY204) (BD Bioscience), NF-κB p65 (pS536) (Cell Signaling Technologies) or isotype control IgG1κ (BD Biosciences) for 30 min at RT. After washing data was acquired on an LSR Fortessa (BD Biosciences) and analyzed in with FlowJo v10.0.8 (TreeStar).

### Ca^2+^ Flux Assay

Freshly isolated NK cells were incubated with Fluo-4 for 30 min at 37C in PBS+2%FCS at the recommended dilution (Fluo-4 Imaging kit, Molecular Probes). Cells were then washed twice and incubated with biotinylated CD16 or biotinylated DNAM-1/2B4 (Miltenyi), with the addition of labeled specific antibodies for CD56, CD57, NKG2A, KIR2DL1, KIR2DL1/S1, KIR3DL1/S1 and KIR2DL2/L3/S2, for 10 min at room temperature. The cells were washed once more and placed on ice until assayed. Prior to FACS analysis, the cells were pre-warmed at 37C for 5 min in the presence or absence of GPN (50μM final concentration) or vacuolin-1 (10μM final concentration). Cells were immediately run on FACS for 30s, followed by addition of 10ug/mL Streptavidin and run for a further 4 min. Ionomycin was added at 4uM final concentration and run for a further 1 min. Ca^2+^-flux kinetics were analysed by FlowJo V10.0.8 (TreeStar).

### Retroviral transduction of NK cell lines

Full length human KIR2DL1 and KIR2DL3 were synthesized using standard gene synthesis with codon optimization (Eurofins Genomics, Ebersberg, Germany), and subsequently subcloned into pMSCV using NotI and BamHI cloning sites. The construct for GCAMP6s-NKG7 was synthesized by combining full length GCAMP6s (Chen et al., 2013), with full length NKG7 using a medium 6AA linker sequence (ggaggttcaggaggcagc), and subcloned in pMSCV. Retroviral particles were produced by transfection of Phoenix-ampho 293T cells using Lipofectimine 3000 (Life Technologies). YTS cells were spinoculated with viral supernatants for 60mins at 1000xG. Cells were screened at 5 passages and positive cells were sorted using a FacsARIA.

### siRNA Interference

Primary NK cells were isolated, rested for 2 hours and transfected either directly or primed with 10ng/mL IL15 for 72 hours and transfected. NK cells or cell lines were transfected by Amaxa nucleofection (Lonza) using 300pM of Dharmacon ON-TARGET plus *SMART*pool control siRNA, or *SMART*pool RNA targeting human TRPML1. Nuclefection was performed using the human macrophage kit using program Y-010. After nucleofection, cells were rested for 4 hours in OPTI-MEM, before an equal volume of culture medium with 2ng/ml IL15 was added. The cells were then cultured for 48 hours before phenotypic and functional testing. siRNA efficiency was determined using qPCR.

### Cytotoxicity assay by Flow Cytometry

Target cell killing was determined using a combination of viability stains. Cytotoxicity assays were performed using an NK cell to target cell ratio of 5:1 at 37C for 5 hours, after which cells were stained surface markers (CD56) to discriminate NK cells from target cells, with the addition of Live/dead aqua fluorescent reactive dye (1:200; LifeTechnologies) for 20 min at 4°C (or 15 min at RT), washed in staining buffer and stained in 50μl RPMI media plus 1μM Yo-Pro^®^-3 iodide (LifeTechnologies) for 15’ at 37°C. Finally, cells were washed and either directly analysed using a BD™ LSR-II cytometer or fixed in 100µl PFA 2-4% for 10 min at 4°C, pelleted, washed twice with 200µl staining buffer and then rested at 4 °C until analysed at the LSR-II machine.

### RNA Sequencing of Education Subsets

RNA was isolated from sorted KIR single positive NK cell subsets. Library preparation was performed using the Illumina NeoPrep Library preparation system. Sequencing was performed using the NextSeq (Illumina) (single read, 75base pairs). Read alignment was carried out using Bowtie (version 2.0.5.0) and Tophat (version 2.0.6), and transcript abundance was estimated using Cufflinks (version 2.1.1). The resulting FPKM values for each transcript were log2 transformed for visualization in scatterplots (Figure 2D).

### Quantitative PCR

RNA was isolated using RNeasy mini kit (Qiagen). Following RNA isolation, cDNA was synthesized using First strand synthesis kit (Qiagen) according to manufacturer’s protocol. Customized RT2 Profiler PCR array (Qiagen) was ordered with specific primers for the 20 genes of interest, as well as 2 housekeeping genes, a reverse transcriptase control, genomic DNA control, and a positive PCR control. Real time quantitative PCR was performed on cDNA from differentiation and education subsets and the data obtained was normalized using 18S rRNA and B2M as housekeeping genes. All target genes were run as triplicates and analysis of qPCR data was done using qBase+ (Biogazelle).

### Statistical analysis

Comparisons of matched groups were made using paired Students T test. Single comparison of groups or populations of cells between donors was performed using Students t test or Mann-Whitney test for statistical significance. n.s. indicates not significant; ****p < 0.0001***; p < 0.001; **p < 0.01; and *p < 0.05. Analyses were performed using GraphPad Prism software.

## Author contributions

JPG designed and performed research, analysed data and wrote the paper. BJ, DC, ES and AB performed imaging experiments and analysed data. LM-Z, SL, TC, UK, SL and MLS performed RNA Seq and qPCR and analysed data. WEL contributed to the Ca-signaling experiments. JL performed Phospho-flow experiments. AP, MTW, EH A, LL and VSO performed experiments. SP, CG, KT and HS contributed to the design of research and the writing of the paper. KJM designed research, analysed data and wrote the paper.

## Acknowledgement

This work was supported by grants from the Swedish Research Council, the Swedish Children’s Cancer Society, the Swedish Cancer Society, the Tobias Foundation, the Karolinska Institutet, the Wenner-Gren Foundation, the Norwegian Cancer Society, the Norwegian Research Council, the South-Eastern Norway Regional Health Authority and the KG Jebsen Center for Cancer Immunotherapy. SP was supported by BB/N01524X/1 from the BBSRC and BJ was funded by a Mildred Scheel postdoctoral scholarship from the Dr. Mildred Scheel Foundation for Cancer Research of the German Cancer Aid Organization.

## References

1. Karre K., Ljunggren H.G., Piontek G. & Kiessling R. Selective rejection of H-2-deficient lymphoma variants suggests alternative immune defence strategy. Nature 319, 675–678 (1986).

2. Anfossi N. et al. Human NK cell education by inhibitory receptors for MHC class I. Immunity 25, 331–342 (2006).

3. Elliott J.M. & Yokoyama W.M. Unifying concepts of MHC-dependent natural killer cell education. Trends Immunol 32, 364–372 (2011).

4. Kim S. et al. Licensing of natural killer cells by host major histocompatibility complex class I molecules. Nature 436, 709–713 (2005).

5. Ebihara T., Jonsson A.H. & Yokoyama W.M. Natural killer cell licensing in mice with inducible expression of MHC class I. Proc Natl Acad Sci U S A 110, E4232–4237 (2013).

6. Jonsson A.H., Yang L., Kim S., Taffner S.M. & Yokoyama W.M. Effects of MHC class I alleles on licensing of Ly49A+ NK cells. J Immunol 184, 3424–3432 (2010).

7. Jonsson A.H. & Yokoyama W.M. Assessing licensing of NK cells. Methods Mol Biol 612, 39–49 (2010).

8. Bjorkstrom N.K. et al. Expression patterns of NKG2A, KIR, and CD57 define a process of CD56dim NK-cell differentiation uncoupled from NK-cell education. Blood 116, 3853–3864 (2010).

9. Brodin P., Lakshmikanth T., Johansson S., Karre K. & Hoglund P. The strength of inhibitory input during education quantitatively tunes the functional responsiveness of individual natural killer cells. Blood (2008).

10. Chen S. et al. The Self-Specific Activation Receptor SLAM Family Is Critical for NK Cell Education. Immunity 45, 292–304 (2016).

11. Joncker N.T., Shifrin N., Delebecque F. & Raulet D.H. Mature natural killer cells reset their responsiveness when exposed to an altered MHC environment. J Exp Med 207, 2065–2072 (2010).

12. Elliott J.M., Wahle J.A. & Yokoyama W.M. MHC class I-deficient natural killer cells acquire a licensed phenotype after transfer into an MHC class I-sufficient environment. J Exp Med 207, 2073–2079 (2010).

13. Viant C. et al. SHP-1-mediated inhibitory signals promote responsiveness and anti-tumour functions of natural killer cells. Nat Commun 5, 5108 (2014).

14. Guia S. et al. Confinement of activating receptors at the plasma membrane controls natural killer cell tolerance. Science signaling 4, ra21 (2011).

15. Staaf E. et al. Educated natural killer cells show dynamic movement of the activating receptor NKp46 and confinement of the inhibitory receptor Ly49A. Science signaling 11 (2018).

16. Enqvist M. et al. Coordinated Expression of DNAM-1 and LFA-1 in Educated NK Cells. J Immunol 194, 4518–4527 (2015).

17. van Bergen, J. et al. HLA reduces killer cell Ig-like receptor expression level and frequency in a humanized mouse model. J Immunol 190, 2880–2885 (2013).

18. Boudreau J.E. & Hsu K.C. Natural killer cell education in human health and disease. Curr Opin Immunol 50, 102–111 (2018).

19. Brodin P. & Hoglund P. Beyond licensing and disarming: a quantitative view on NK-cell education. Eur J Immunol 38, 2934–2937 (2008).

20. Valiante N.M. et al. Functionally and structurally distinct NK cell receptor repertoires in the peripheral blood of two human donors. Immunity 7, 739–751 (1997).

21. Andersson S., Fauriat C., Malmberg J.A., Ljunggren H.G. & Malmberg K.J. KIR acquisition probabilities are independent of self-HLA class I ligands and increase with cellular KIR expression. Blood 114, 95–104 (2009).

22. Lopez-Verges S. et al. CD57 defines a functionally distinct population of mature NK cells in the human CD56dimCD16+ NK-cell subset. Blood 116, 3865–3874 (2010).

23. Boudreau J.E., Mulrooney T.J., Le Luduec J.B., Barker E. & Hsu K.C. KIR3DL1 and HLA-B Density and Binding Calibrate NK Education and Response to HIV. J Immunol 196, 3398–3410 (2016).

24. Fauriat C. et al. Estimation of the size of the alloreactive NK cell repertoire: studies in individuals homozygous for the group A KIR haplotype. J Immunol 181, 6010–6019 (2008).

25. Yawata M. et al. MHC class I-specific inhibitory receptors and their ligands structure diverse human NK-cell repertoires toward a balance of missing self-response. Blood 112, 2369–2380 (2008).

26. Yu J. et al. Hierarchy of the human natural killer cell response is determined by class and quantity of inhibitory receptors for self-HLA-B and HLA-C ligands. J Immunol 179, 5977–5989 (2007).

27. Bryceson Y.T. et al. Molecular mechanisms of natural killer cell activation. Journal of innate immunity 3, 216–226 (2011).

28. Fehniger T.A. et al. Acquisition of murine NK cell cytotoxicity requires the translation of a pre-existing pool of granzyme B and perforin mRNAs. Immunity 26, 798–811 (2007).

29. Mace E.M. et al. NK cell lytic granules are highly motile at the immunological synapse and require F-actin for post-degranulation persistence. J Immunol 189, 4870–4880 (2012).

30. Sutton V.R. et al. Serglycin determines secretory granule repertoire and regulates natural killer cell and cytotoxic T lymphocyte cytotoxicity. FEBS J 283, 947–961 (2016).

31. Kolset S.O. & Pejler G. Serglycin: a structural and functional chameleon with wide impact on immune cells. J Immunol 187, 4927–4933 (2011).

32. Gwalani L.A. & Orange J.S. Single Degranulations in NK Cells Can Mediate Target Cell Killing. J Immunol 200, 3231–3243 (2018).

33. Forslund E. et al. Microchip-Based Single-Cell Imaging Reveals That CD56dimCD57-KIR-NKG2A+ NK Cells Have More Dynamic Migration Associated with Increased Target Cell Conjugation and Probability of Killing Compared to CD56dimCD57-KIR-NKG2A- NK Cells. J Immunol 195, 3374–3381 (2015).

34. Thomas L.M., Peterson M.E. & Long E.O. Cutting edge: NK cell licensing modulates adhesion to target cells. J Immunol 191, 3981–3985 (2013).

35. Davis L.C. et al. NAADP activates two-pore channels on T cell cytolytic granules to stimulate exocytosis and killing. Curr Biol 22, 2331–2337 (2012).

36. Patel S. & Docampo R. Acidic calcium stores open for business: expanding the potential for intracellular Ca2+ signaling. Trends Cell Biol 20, 277–286 (2010).

37. Patel S. & Cai X. Evolution of acidic Ca(2)(+) stores and their resident Ca(2)(+)-permeable channels. Cell Calcium 57, 222–230 (2015).

38. Wolf I.M. et al. Frontrunners of T cell activation: Initial, localized Ca2+ signals mediated by NAADP and the type 1 ryanodine receptor. Sci Signal 8, ra102 (2015).

39. Speak A.O. et al. Altered distribution and function of natural killer cells in murine and human Niemann-Pick disease type C1. Blood 123, 51–60 (2014).

40. Penny C.J., Kilpatrick B.S., Han J.M., Sneyd J. & Patel S. A computational model of lysosome-ER Ca2+ microdomains. J Cell Sci 127, 2934–2943 (2014).

41. Jadot M., Colmant C., Wattiaux-De Coninck S. & Wattiaux R. Intralysosomal hydrolysis of glycyl-L-phenylalanine 2-naphthylamide. Biochem J 219, 965–970 (1984).

42. Sukhai M.A. et al. Lysosomal disruption preferentially targets acute myeloid leukemia cells and progenitors. J Clin Invest 123, 315–328 (2013).

43. Paivandy A. et al. Mefloquine, an anti-malaria agent, causes reactive oxygen species-dependent cell death in mast cells via a secretory granule-mediated pathway. Pharmacol Res Perspect 2, e00066 (2014).

44. Saftig P. & Klumperman J. Lysosome biogenesis and lysosomal membrane proteins: trafficking meets function. Nature reviews. Molecular cell biology 10, 623–635 (2009).

45. Luzio J.P., Pryor P.R. & Bright N.A. Lysosomes: fusion and function. Nature reviews. Molecular cell biology 8, 622–632 (2007).

46. Cao Q., Yang Y., Zhong X.Z. & Dong X.P. The lysosomal Ca(2+) release channel TRPML1 regulates lysosome size by activating calmodulin. J Biol Chem 292, 8424–8435 (2017).

47. Pryor P.R., Reimann F., Gribble F.M. & Luzio J.P. Mucolipin-1 is a lysosomal membrane protein required for intracellular lactosylceramide traffic. Traffic 7, 1388–1398 (2006).

48. Thompson E.G., Schaheen L., Dang H. & Fares H. Lysosomal trafficking functions of mucolipin-1 in murine macrophages. BMC Cell Biol 8, 54 (2007).

49. Dong X.P. et al. PI(3,5)P(2) controls membrane trafficking by direct activation of mucolipin Ca(2+) release channels in the endolysosome. Nat Commun 1,38 (2010).

50. Park S. et al. Fusion of lysosomes with secretory organelles leads to uncontrolled exocytosis in the lysosomal storage disease mucolipidosis type IV. EMBO Rep 17, 266–278 (2016).

51. Sano O. et al. Vacuolin-1 inhibits autophagy by impairing lysosomal maturation via PIKfyve inhibition. FEBS Lett 590, 1576–1585 (2016).

52. Grimm C., Bartel K., Vollmar A.M. & Biel M. Endolysosomal Cation Channels and Cancer-A Link with Great Potential. Pharmaceuticals (Basel) 11 (2018).

53. Chen C.C. et al. A small molecule restores function to TRPML1 mutant isoforms responsible for mucolipidosis type IV. Nat Commun 5, 4681 (2014).

54. Shen D. et al. Lipid storage disorders block lysosomal trafficking by inhibiting a TRP channel and lysosomal calcium release. Nat Commun 3, 731 (2012).

55. Raulet D.H. & Vance R.E. Self-tolerance of natural killer cells. Nat Rev Immunol 6, 520–531 (2006).

56. Er E.E., Mendoza M.C., Mackey A.M., Rameh L.E. & Blenis J. AKT facilitates EGFR trafficking and degradation by phosphorylating and activating PIKfyve. Science signaling 6, ra45 (2013).

57. Bretou M. et al. Lysosome signaling controls the migration of dendritic cells. Sci Immunol 2 (2017).

58. Hockey L.N. et al. Dysregulation of lysosomal morphology by pathogenic LRRK2 is corrected by TPC2 inhibition. J Cell Sci 128, 232–238 (2015).

59. Wagner J.A. et al. CD56bright NK cells exhibit potent antitumor responses following IL-15 priming. J Clin Invest 127, 4042–4058 (2017).

60. Thompson T.W. et al. Endothelial cells express NKG2D ligands and desensitize anti-tumor NK responses. Elife 6 (2017).

61. Kwon H.J. et al. Stepwise phosphorylation of p65 promotes NF-kappaB activation and NK cell responses during target cell recognition. Nat Commun 7, 11686 (2016).

62. Upshaw J.L. et al. NKG2D-mediated signaling requires a DAP10-bound Grb2-Vav1 intermediate and phosphatidylinositol-3-kinase in human natural killer cells. Nat Immunol 7, 524–532 (2006).

63. Lodeiro M. et al. The SHP-1 protein tyrosine phosphatase negatively modulates Akt signaling in the ghrelin/GHSR1a system. Mol Biol Cell 22, 4182–4191 (2011).

64. Bryceson Y.T. & Long E.O. Line of attack: NK cell specificity and integration of signals. Curr Opin Immunol 20, 344–352 (2008).

65. Stebbins C.C. et al. Vav1 dephosphorylation by the tyrosine phosphatase SHP-1 as a mechanism for inhibition of cellular cytotoxicity. Mol Cell Biol 23, 6291–6299 (2003).

66. Gumbleton M. et al. Dual enhancement of T and NK cell function by pulsatile inhibition of SHIP1 improves antitumor immunity and survival. Science signaling 10 (2017).

67. Gumbleton M., Vivier E. & Kerr W.G. SHIP1 intrinsically regulates NK cell signaling and education, resulting in tolerance of an MHC class I-mismatched bone marrow graft in mice. J Immunol 194, 2847–2854 (2015).

68. Chen C.S., Bach G. & Pagano R.E. Abnormal transport along the lysosomal pathway in mucolipidosis, type IV disease. Proc Natl Acad Sci U S A 95, 6373–6378 (1998).

69. Miller A. et al. Mucolipidosis type IV protein TRPML1-dependent lysosome formation. Traffic 16, 284–297 (2015).

70. Shaik G.M., Draberova L., Heneberg P. & Draber P. Vacuolin-1-modulated exocytosis and cell resealing in mast cells. Cell Signal 21, 1337–1345 (2009).

71 . Choy C.H. et al. Lysosome enlargement during inhibition of the lipid kinase PIKfyve proceeds through lysosome coalescence. J Cell Sci (2018).

72. Nagle D.L. et al. Identification and mutation analysis of the complete gene for Chediak-Higashi syndrome. Nat Genet 14, 307–311 (1996).

73. Gil-Krzewska A. et al. Chediak-Higashi syndrome: Lysosomal trafficking regulator domains regulate exocytosis of lytic granules but not cytokine secretion by natural killer cells. J Allergy Clin Immunol 137, 1165–1177 (2016).

74. Chiang, S.C.C. et al. Differences in Granule Morphology yet Equally Impaired Exocytosis among Cytotoxic T Cells and NK Cells from Chediak-Higashi Syndrome Patients. Frontiers in immunology 8, 426 (2017).

75. Gil-Krzewska A. et al. An actin cytoskeletal barrier inhibits lytic granule release from Natural Killer cells in Chediak-Higashi syndrome. J Allergy Clin Immunol (2017).

76. Chen H.Y. et al. Nanoimaging granule dynamics and subcellular structures in activated mast cells using soft X-ray tomography. Sci Rep 6, 34879 (2016).

77. Isaaz S., Baetz K., Olsen K., Podack E. & Griffiths G.M. Serial killing by cytotoxic T lymphocytes: T cell receptor triggers degranulation, re-filling of the lytic granules and secretion of lytic proteins via a non-granule pathway. Eur J Immunol 25, 1071–1079 (1995).

78. Goodridge J.P., Onfelt B. & Malmberg K.J. Newtonian cell interactions shape natural killer cell education. Immunol Rev 267, 197–213 (2015).

79. Landtwing V. et al. Cognate HLA absence in trans diminishes human NK cell education. J Clin Invest (2016).

80. Boudreau J.E. et al. Cell-Extrinsic MHC Class I Molecule Engagement Augments Human NK Cell Education Programmed by Cell-Intrinsic MHC Class I. Immunity 45, 280–291 (2016).

81. Orange J.S. et al. Wiskott-Aldrich syndrome protein is required for NK cell cytotoxicity and colocalizes with actin to NK cell-activating immunologic synapses. Proc Natl Acad Sci U S A 99, 11351–11356 (2002).

82. Anton O.M., Vielkind S., Peterson M.E., Tagaya Y. & Long E.O. NK Cell Proliferation Induced by IL-15 Transpresentation Is Negatively Regulated by Inhibitory Receptors. J Immunol 195, 4810–4821 (2015).

83. Marcais A. et al. The metabolic checkpoint kinase mTOR is essential for IL-15 signaling during the development and activation of NK cells. Nat Immunol 15, 749–757 (2014).

84. Marcais A. et al. High mTOR activity is a hallmark of reactive natural killer cells and amplifies early signaling through activating receptors. Elife 6 (2017).

85. Zoncu R. et al. mTORC1 senses lysosomal amino acids through an inside-out mechanism that requires the vacuolar H(+)-ATPase. Science 334,678–683 (2011).

